# Association of homelessness and diet on the gut microbiome: A United States-Veteran Microbiome Project (US-VMP) study

**DOI:** 10.1101/2023.09.22.559004

**Authors:** Andrew J. Hoisington, Kelly A. Stearns-Yoder, Christopher E. Stamper, Ryan Holliday, Diana P. Brostow, Molly E. Penzenik, Jeri E. Forster, Teodor T. Postolache, Christopher A. Lowry, Lisa A. Brenner

**Author notes:** Corresponding author Lisa A. Brenner, Ph.D.

## Abstract

Military Veterans account for 8% of homeless individuals living in the United States. To highlight associations between history of homelessness and the gut microbiome, we compared the gut microbiome of Veterans who reported having a previous experience of homelessness to those from individuals who reported never having experienced a period of homelessness. Moreover, we examined the impact of the cumulative exposure of prior and current homelessness to understand possible associations between these experiences and the gut microbiome. Microbiome samples underwent genomic sequencing and were analyzed based on alpha diversity, beta diversity, and taxonomic differences. Additionally, demographic information, dietary data, and mental health history were collected. A lifetime history of homelessness was found to be associated with alcohol use disorder, substance use disorder, and healthy eating index compared to those without such a history. In terms of differences in gut microbiota, beta diversity was significantly different between Veterans that had experienced homelessness and Veterans that had never been homeless (*p* = 0.047, Weighted UniFrac), while alpha diversity was similar. The microbial community differences were, in part, driven by a lower relative abundance of *Akkermansia* in Veterans that had experienced homelessness (mean ± SD; 1.07 ± 3.85) compared to Veterans that had never been homeless (2.02 ± 5.36) (*p* = 0.014, ancom-bc2). Additional research is required to facilitate understanding regarding complex associations between homelessness, the gut microbiome, and mental and physical health conditions, with a focus on increasing understanding regarding the longitudinal impact of housing instability throughout the lifespan.

**Importance:** Although there are known stressors related to homelessness, as well as chronic health conditions experienced by those without stable housing, there has been limited work evaluating the associations between microbial community composition and homelessness. We analyzed, for the first time, bacterial gut microbiome associations among those with experiences of homelessness on alpha diversity, beta diversity, and taxonomic differences. Additionally, we characterized the influences of diet, demographic characteristics, military service history and mental health conditions on the microbiome of Veterans with and without any lifetime history of homelessness. Future longitudinal research to evaluate the complex relationships between homelessness, the gut microbiome, and mental health outcomes is recommended. Ultimately, differences in the gut microbiome of individuals experiencing and not experiencing homelessness could assist in identification of treatment targets to improve health outcomes.

## Introduction

Homelessness can be defined as lacking a: “fixed, regular, and adequate nighttime residence, such as … living in emergency shelters, transitional housing, or places not meant for habitation” (1). Addressing homelessness among Veterans remains a top priority for the United States (US) Department of Veterans Affairs (VA). Nonetheless, on a single night in 2020, 37,252 Veterans were noted to be experiencing homelessness (2). This accounts for approximately 8% of all homeless adults in the US (2). Among homeless Veterans surveyed, over 40% were determined to be unsheltered (e.g., sleeping outside or in their car) (2), with the remainder residing in sheltered locations such as emergency shelters or transitional housing programs. Additionally, some forms of homelessness (e.g., “couch-surfing”) are frequently undercounted (3), especially among women (4).

Veterans experiencing homelessness experience a wide range of physical and mental health conditions (5, 6), and a lack of housing is particularly associated with substance misuse, death by suicide, and infectious- (e.g., tuberculosis) and injury-related health outcomes (e.g., traumatic brain injury) (7–9). Moreover, Veterans experiencing homelessness often have psychosocial challenges (e.g., unemployment, criminal legal involvement), which can further negatively impact both their ability to access resources (e.g., stable housing, medical care, food (10)), and their overall health (e.g., high rates of metabolic conditions among those with serious and persistent mental illness) (11, 12). For example, Veterans that are homeless are prone to experiencing food insecurity, which may exacerbate malnutrition and poor health (13). In combination, these challenges have been noted to result in chronic stress (14), which has been associated with enduring physical and mental health conditions (e.g., cardiovascular disease, gastrointestinal disorders, depression, and substance use) (8, 15).

Chronic stress is associated with altered autonomic nervous system activity, impaired regulation of the hypothalamic-pituitary-adrenal (HPA) axis, impaired glucocorticoid-signaling, and a systemic low-grade proinflammatory state, as well as adverse physical and psychological outcomes (16–20). An area of relatively recent research includes the exploration of associations between the human microbiome, inflammation, and health outcomes (21). Microbial colonization of the gut has been shown to influence the development of the immune system (22).

Moreover, an imbalance or depletion of the intestinal ecosystem may alter immune responses (23), and contribute to systemic physiologic dysfunction, including elevated inflammation and oxidative stress (24–26). Researchers have also suggested significant associations between microbiome and chronic health conditions (27–29). For example, homeless individuals experience an elevated risk of hepatitis C viral infection (30) which in turn can have long-term effects on gut microbiome composition, even after viral eradication (31).

Nonetheless, despite known stressors related to homelessness, as well as chronic health conditions experienced by those without stable housing, there has been limited work evaluating associations between microbial community composition and homelessness. As such, this study aimed to compare the gut microbiome from Veterans who reported having a previous experience of homelessness (lifetime) to those from individuals who never experienced any period of homelessness. Moreover, we further examined the impact of cumulative exposures of homelessness (number of experiences) and current homelessness to understand these experiences on the Veteran gut microbiome.

## Materials and Methods

### Sample Description

The United States-Veteran Microbiome Project (US-VMP) has previously been described Although the US-VMP is longitudinal in nature, only baseline (Time 1) data collected from 2016-2020 were used in this study. We compared gut microbial samples from Veterans that reported having a lifetime experience of homelessness (*n* = 160) to those who had no reported history of homelessness (*n* = 228). Further exploratory analyses were conducted regarding the influence of the total number of reported experiences of homelessness on microbial composition, as well as current homelessness. This study was conducted according to the guidelines of the Declaration of Helsinki and all procedures involving human participants were approved by the Colorado Multiple Institutional Review Board (COMIRB #15-1885), as well as the local VA regulatory committee. Written informed consent was obtained before Veterans participated in any study procedures. All participants were US Veterans that were seeking VA care.

### Measures

For a comprehensive list of measures administered in the US-VMP study, see Brenner et al. (33). Data from the measures listed in Tables in the manuscript (**Table 1** and **Table 2)** and and supplemental files (**Tables S1 & S2)** were included in these analyses. The Rocky Mountain MIRECC Demographics Questionnaire includes standard demographic questions and information regarding histories of military service and homelessness. Three homelessness-related questions were presented to participants that queried as to whether individuals were currently homeless, or had a lifetime history of homelessness, as well as the number of unique periods of homelessness they had experienced. The participants were not provided a definition of homelessness by the study team.

**Table 1.** Demographic Characteristics for Veterans in the United States-Veteran Microbiome Project by lifetime history of homelessness.

|  | Full Sample | No Lifetime History of Homelessness | Lifetime History of Homelessness | *p* for difference | Test used |
| --- | --- | --- | --- | --- | --- |
| Variable | *N* (%) or Mean ± SD (range) | *N* (%) or Mean ± SD (range) | *N* (%) or Mean ± SD (range) |  |  |
| Total | *N* = 388 | *N* = 228 | *N* = 160 |  |  |
| Age | 48.2 ± 13.2 (23-85) | 45.7 ± 13.8 (23-85) | 51.9 ± 11.3 (25-76) | <0.0001 | Independent samples *t*-test |
| Age Categories | | | | <0.0001 | $\chi^2$ |
| 20-29 | 28 (7.2%) | 21 (9.2%) | 7 (4.4%) |  |  |
| 30-39 | 97 (25.0%) | 73 (32.0%) | 24 (15.0%) |  |  |
| 40-49 | 74 (19.1%) | 50 (21.9%) | 24 (15.0%) |  |  |
| 50-59 | 98 (25.3%) | 38 (16.7%) | 60 (37.5%) |  |  |
| 60-69 | 74 (19.1%) | 33 (14.5%) | 41 (25.6%) |  |  |
| 70+ | 17 (4.4%) | 13 (5.7%) | 4 (2.5%) |  |  |
| Gender<sup>1</sup> |  |  |  | 0.01 | Fisher's exact test |
| Male | 311 (80.4%) | 174 (76.7%) | 137 (85.6%) |  |  |
| Female | 74 (19.1%) | 53 (23.4%) | 21 (13.1%) |  |  |
| Transgender | 1 (0.3%) | 0 | 1 (0.6%) |  |  |
| Intersex | 1 (0.3%) | 0 | 1 (0.6%) |  |  |
| Race<sup>2</sup> |  |  |  | <0.0001 | Fisher's exact test |
| Caucasian | 261 (68.3%) | 179 (79.6%) | 82 (52.2%) |  |  |
| African American | 65 (17.0%) | 24 (10.7%) | 41 (26.1%) |  |  |
| Native American/<br>Alaskan Native | 6 (1.6%) | 2 (0.9%) | 4 (2.6%) |  |  |
| Asian | 3 (0.8%) | 3 (1.3%) | 0 |  |  |
| Pacific Islander | 1 (0.3%) | 1 (0.4%) | 0 |  |  |
| Multiracial | 22 (5.8%) | 8 (3.6%) | 14 (8.9%) |  |  |
| Other | 24 (6.3%) | 8 (3.6%) | 16 (10.2%) |  |  |
| Ethnicity<sup>2</sup> | | | | 0.13 | $\chi^2$ |
| Hispanic | 58 (15.2%) | 29 (12.9%) | 29 (18.5%) |  |  |
| Non-Hispanic | 324 (84.8%) | 196 (87.1%) | 128 (81.5%) |  |  |
| Marital Status<sup>3</sup> | | | | <0.0001 | $\chi^2$ |
| Married | 154 (40.0%) | 124 (54.9%) | 30 (18.9%) |  |  |
| Single | 90 (23.4%) | 46 (20.4%) | 44 (27.7%) |  |  |
| Cohabiting | 21 (5.5%) | 14 (6.2%) | 7 (4.4%) |  |  |
| Separated/Divorced | 105 (27.3%) | 36 (15.9%) | 69 (43.4%) |  |  |
| Widowed | 15 (3.9%) | 6 (2.7%) | 9 (5.7%) |  |  |
| Sexual Orientation<sup>4</sup> |  |  |  | 0.03 | Fisher's exact test |
| Heterosexual | 353 (91.5%) | 208 (92.0%) | 145 (90.6%) |  |  |
| Gay/Lesbian/Queer | 20 (5.2%) | 9 (4.0%) | 11 (6.9%) |  |  |
| Bisexual | 9 (2.3%) | 8 (3.5%) | 1 (0.6%) |  |  |
| Questioning | 1 (0.3%) | 1 (0.4%) | 0 |  |  |
| Other | 3 (0.8%) | 0 | 3 (1.9%) |  |  |

| Education Level<sup>1</sup> |  |  |  |  |  |
| --- | --- | --- | --- | --- | --- |
| No High School Degree | 1 (0.3%) | 0 | 1 (0.6%) | <0.0001 | Fisher's exact test |
| High School Degree | 41 (10.6%) | 9 (4.0%) | 32 (20.0%) |  |  |
| Some College | 133 (34.4%) | 68 (30.0%) | 65 (40.6%) |  |  |
| Associate Degree | 66 (17.1%) | 39 (17.2%) | 27 (16.9%) |  |  |
| Bachelor Degree | 89 (23.0%) | 64 (28.2%) | 25 (15.6%) |  |  |
| Master's Degree | 50 (12.9%) | 41 (18.1%) | 9 (5.6%) |  |  |
| Doctoral Degree | 7 (1.8%) | 6 (2.6%) | 1 (0.6%) |  |  |
| Employment Status<sup>5</sup> |  |  |  |  |  |
| Employed Full-Time | 95 (24.8%) | 71 (31.7%) | 24 (15.1%) | <0.0001 | $\chi^2$ |
| Employed Part-Time | 40 (10.4%) | 28 (12.5%) | 12 (7.6%) |  |  |
| Unemployed, Seeking Job | 59 (15.4%) | 24 (10.7%) | 35 (22.0%) |  |  |
| Unemployed, Not Seeking Job | 77 (20.1%) | 37 (16.5%) | 40 (25.2%) |  |  |
| Retired | 112 (29.2%) | 64 (28.6%) | 48 (30.2%) |  |  |
| Student Status<sup>1</sup> |  |  |  |  |  |
| Not in school | 324 (83.7%) | 175 (76.8%) | 149 (93.7%) | <0.0001 | $\chi^2$ |
| Part-Time | 16 (4.1%) | 13 (5.7%) | 3 (1.9%) |  |  |
| Full-Time | 47 (12.1%) | 40 (17.5%) | 7 (4.4%) |  |  |
| Currently Homeless<sup>4</sup> |  |  |  |  |  |
| Yes | 31 (8.0%) | 0 | 31 (19.6%) | <0.0001 | $\chi^2$ |
| No | 355 (92.0%) | 228 (100.0%) | 127 (80.4%) |  |  |
| Number of Times Ever Homeless | 1.5 ± 5.6 (0-100) | 0 | 3.6 ± 8.3 (1-100) | <0.0001 | Wilcoxon rank-sum test |

Note: *p* value and test between no lifetime history of homelessness and lifetime history of homelessness; <sup>1</sup>1 Missing, <sup>2</sup>6 Missing, <sup>3</sup>3 Missing, <sup>4</sup>2 Missing, <sup>5</sup>5 Missing

**Table 2.** Total Healthy Eating Index (HEI) and component indices for Veterans in the United States-Veteran Microbiome Project by lifetime history of homelessness.

|  | **Full Sample**<br>**N = 387** | **No History of Homelessness**<br>**N = 228** | **Lifetime History of Homelessness**<br>**N = 159\*** | **p for difference** | **Test used** |
| --- | --- | --- | --- | --- | --- |
| | Mean ± SD | Mean ± SD | Mean ± SD | | |
| Total HEI | 66.9 ± 11.0 | 68.9 ± 10.2 | 64.0 ± 11.5 | **<.0001** | Independent samples *t*-test |
| **HEI Sub-scores** |  |  |  |  |  |
| Total Vegetables | 4.2 ± 1.2 | 4.3 ± 1.2 | 4.1 ± 1.3 | 0.10 | Independent samples *t*-test |
| Beans & Greens | 4.3 ± 1.3 | 4.4 ± 1.2 | 4.2 ± 1.3 | 0.21 | Independent samples *t*-test |
| Total Fruits | 3.7 ± 1.5 | 3.9 ± 1.5 | 3.5 ± 1.5 | **0.03** | Independent samples *t*-test |
| Whole Fruits | 4.2 ± 1.4 | 4.3 ± 1.3 | 4.0 ± 1.4 | 0.11 | Independent samples *t*-test |
| Total Protein Foods | 2.9 ± 0.7 | 3.0 ± 0.7 | 2.8 ± 0.6 | **0.0009** | Independent samples *t*-test |
| Seafood & Plant Proteins | 4.3 ± 0.9 | 4.4 ± 0.8 | 4.2 ± 0.9 | **0.03** | Independent samples *t*-test |
| Whole Grains | 4.1 ± 2.2 | 4.3 ± 2.3 | 3.9 ± 2.1 | **0.05** | Independent samples *t*-test |
| Dairy | 3.2 ± 2.8 | 3.1 ± 2.8 | 3.2 ± 2.8 | 0.89 | Independent samples *t*-test |
| Fatty Acids | 4.6 ± 2.9 | 5.0 ± 2.9 | 3.9 ± 2.8 | **<.0001** | Independent samples *t*-test |
| Sodium | 8.9 ± 1.6 | 9.0 ± 1.6 | 8.9 ± 1.7 | 0.35 | Independent samples *t*-test |
| Saturated Fats | 5.5 ± 2.8 | 5.7 ± 2.9 | 5.2 ± 2.7 | 0.07 | Independent samples *t*-test |
| Added Sugars | 7.0 ± 3 | 7.5 ± 2.8 | 6.3 ± 3.3 | **<.0001** | Independent samples *t*-test |
| Refined Grains | 9.9 ± 0.8 | 10.0 ± 0.7 | 9.9 ± 0.9 | 0.16 | Wilcoxon rank-sum test |

Note: *p* value and test between no history of homelessness and lifetime history of homelessness; \*One participant was missing HEI data.

The primary mental health measure was the Structured Clinical Interview for DSM-5-TR Axis I Disorders, Research Version, Patient Edition with Psychotic Screen (SCID-5-I/P W/ PSY SCREEN), which is a comprehensive semi-structured clinical interview for the assessment of major Diagnostic and Statistical Manual of Mental Disorders-5 diagnoses (34). Participants also completed the 36-Item Short Form Health Survey (SF-36) (35), a 36-item survey with scores ranging from 0-100 and subscales regarding perceived physical (Physical Component Summary; SF-36 PCS) and mental (Mental Component Summary; SF-36 MCS) functioning, with higher scores being associated with better perceived functioning.

### Diet

Dietary data were evaluated using the Harvard Willett Food-Frequency Questionnaire (FFQ) (36). The FFQ consists of over one hundred items in 18 categories, and participants are asked to report how often they consumed a standardized portion of each type of food/beverage in the previous year (e.g., ½ cup cooked spinach, 1 banana, 5-ounce glass of wine), with possible responses ranging from “never” to “6 or more times per day”. The Healthy Eating Index (HEI) (37) was applied to evaluate dietary quality. The HEI-2015 was created to assess adherence to the 2015-2020 U.S. Dietary Guidelines for Americans established by the U.S. Departments of Agriculture, and Health and Human Services (38) and can be used to identify 13 different energy-adjusted variables. The HEI yields individual subscales for each variable, and as an aggregate score ranging from 0 to 100, with 0 being no adherence to dietary guidelines, and 100 being perfect adherence.

### Microbiome Sample Collection and Processing

Microbiome sample collection for the US-VMP has been previously described in detail Briefly, sterile dual tipped swabs were used to collect fecal microbiome samples. Samples were collected either in person during the study visit or by the participant at their residence and shipped back to the research facility where they were stored at –80 °C. Sample DNA was extracted from microbiome samples using the PowerSoil DNA extraction kit (Cat. No. 12955-4, Qiagen, Valencia, California). Marker genes in isolated DNA were polymerase chain reaction (PCR)-amplified using GoTaq Master Mix (Cat. No. M5133, Promega, Madison, Wisconsin) and 515 F (5′-GTGCCAGCMGCCGCGGTAA-3′), 806 R (5′-GGACTACHVGGGTWTCTAAT-3′) primer pair (Integrated DNA Technologies, Coralville, Iowa) targeting the V4 hypervariable region of the 16S rRNA gene modified with a unique 12-base sequence identifier for each sample and the Illumina adapter as previously described in Caporaso et al. (39). The thermal cycling program consisted of an initial step at 94 °C for 3 minutes followed by 35 cycles (94 °C for 45 seconds, 55 °C for 1 minute, and 72 °C for 1.5 minutes), and a final extension at 72 °C for 10 minutes. PCRs were run in duplicate and the products from the duplicate reactions were pooled and visualized on an agarose gel to ensure successful amplification. PCR products were cleaned, normalized, and sequenced by the Colorado Anschutz Research Genetics Organization at the University of Colorado Anschutz Medical Campus. Sequencing was performed on an Illumina MiSeq using V2 chemistry and 300 cycle, 2 × 150-bp paired-end sequencing.

### Microbiome Sample Analysis

Sequencing data were processed and analyzed using the Quantitative Insights Into Microbial Ecology program (QIIME2 v. 2021.8). The Deblur algorithm (40) was used to denoise demultiplexed sequences. Phylogenic assignment was augmented through SATe-enabled phylogenetic placement (SEPP (41)) to improve accuracy prior to assignment of taxonomic classification based on the silva_128 database (42). No samples had any outlying amplicon sequencing variants (ASVs) removed from the SEPP process. Samples that were shipped to the research facility had taxa removed that are known to artificially “bloom” during shipping (43).

For α- and β-diversity and taxonomic evaluations, samples were rarefied at 7,000 sequences per sample. After quality control, 20 samples were removed, and the remaining 388 samples were analyzed. Demultiplexed single-end sequences were deposited in the NCBI Sequence Read Archive (BioProject accession ID: PRJNA1010779).

Analyses were performed with the open-source statistical software R v. 3.5.1 (44) (https://www.R-project.org) and QIIME2-2021.8. The α-diversity metrics assessed were (1) Observed features, also called Observed ASVs; (2) Shannon diversity; and (3) Pielou’s Evenness. All statistical tests were conducted with a two-tailed alpha level of 0.05. Comparison between homeless categories for each α-diversity metric was completed in R using the “glm” function for generalized linear modeling and adjusted for age, gender, and education level. The three covariates were not colinear (Pearson Correlation Coefficients -0.01 to 0.17). For β-diversity, principal coordinates analysis (PCoA) was performed using the vegan package (45) for Unweighted UniFrac and Weighted UniFrac (46). Statistical differences for categories of homelessness were calculated through pairwise PERMANOVA with 10,000 permutations through the vegan package (47) with “adonis2” function adjusted for age, gender, and education level. In order to determine the strongest correlates of fecal microbiota composition, we used a stepwise model-building tool for constrained ordination methods based on adjusted *r^2^*, using the UniFrac distance metrics with the “envfit” function of the vegan package that uses PERMANOVA for significance. Canonical correspondence analysis (CCA) was applied to distance metrics for biplot generation and to determine the influence of individual variables on the microbial community. Differential relative abundances of taxa based on homeless categories were assessed using the “ancom-bc2” function (48) with covariates of age, gender, and education level (prevalence filter 30%, *p* adjusted for false discovery rate of multiple tests via Benjamini-Hochberg method (49). Microbial measures of taxonomic relative abundance were aggregated at the phylum and genus levels.

## Results

### Cohort Characteristics

Participant demographics, military characteristics, and mental health data are presented in **Table 1** and **Tables S1 & S2**. Veterans with a lifetime history of homelessness were significantly more likely to be older (mean age ± SD, lifetime history of homelessness, 51.9 ± 11.3 years; never homeless, 45.7 ± 13.8 years), male, and racially diverse when compared to those with no reported homelessness. Higher age was expected in Veterans with a lifetime history of homelessness, due to more time available to become homeless. Members of the cohort with lifetime experience of homelessness were significantly less likely to be married, have advanced degrees, and be employed part- or full-time. They also often noted multiple experiences of homelessness, with 59.4% of Veterans that had a lifetime history of homelessness reporting two or more experiences. Thirty-one individuals were currently homeless at the time of study participation.

The majority of Veterans with a lifetime history of homelessness served in the post-Vietnam/pre-Gulf War era. They had also been discharged from the military (on average) over ten years prior to the study period, when compared to Veterans that did not report a lifetime history of homelessness. Those with a lifetime history of homelessness served significantly less time in the military (considering both Active and Reserve or National Guard duty), and had fewer deployments and periods of service in combat. Regarding lifetime psychiatric disorders, significant differences between those with and without a lifetime history of homelessness were observed for histories of substance use disorder and alcohol use disorder.

### Diet

The mean total HEI score for all participants was 66.9 (Table 2), which is higher than the national average of 59 (50). Veterans with a lifetime history of homelessness scored significantly lower than never homeless counterparts for the mean total HEI score (64.0 compared to 68.9, Independent samples *t*-test, *p* < 0.0001), as well as for Total Fruits, Total Protein Foods, Seafood & Plant Proteins, Whole Grain, Fatty Acids, and Added Sugars (Table 2).

### Fecal Microbiome - Summary

Across all participants (*N* = 388), 3,718 unique amplicon sequencing variants were detected. The taxonomic phyla were dominated by known gut bacterial taxa to include Firmicutes (mean ± SD relative abundance 47.8 ± 21.4) and Bacteroidetes (36.7 ± 22.8). At the genus level, the most abundant taxa were *Bacteroides* (20.6 ± 18.2), *Faecalibacterium* (7.05 ± 7.77), *Blautia* (5.46 ± 6.57), *Prevotella 9* (4.01 ± 10.6), and *Parabacteroides* (3.53 ± 4.08). Participants’ alpha diversity was calculated for Observed ASVs (185 ± 66), Shannon Diversity Index (3.37 ± 0.67), and Pielou’s Evenness (0.65 ± 0.10).

### Fecal Microbiome - Lifetime History of Homelessness

The gut microbiome among those with a lifetime history of homelessness was compared to those individuals who reported that they had never been homeless. There was no difference in alpha diversity, using Observed ASVs, Shannon Diversity Index, and Pielou’s Evenness, between Veterans with and without a lifetime history of homelessness (**Figure S1**). The most abundant genera were similar between Veterans with and without a history of homelessness (**Fig 1**). More specifically, the five most abundant genera for Veterans that experienced homelessness were *Bacteroides, Faecalibacterium, Blautia, Prevotella 9*, and *Parabacteroides.* For Veterans that never experienced homelessness, the first three most abundant genera were consistent with those of Veterans with a lifetime history of homelessness, but *Parabacteroides* was the fourth, and an unnamed genus from *Lachnospiraceae* was the fifth most abundant taxa.

**Fig 1.**
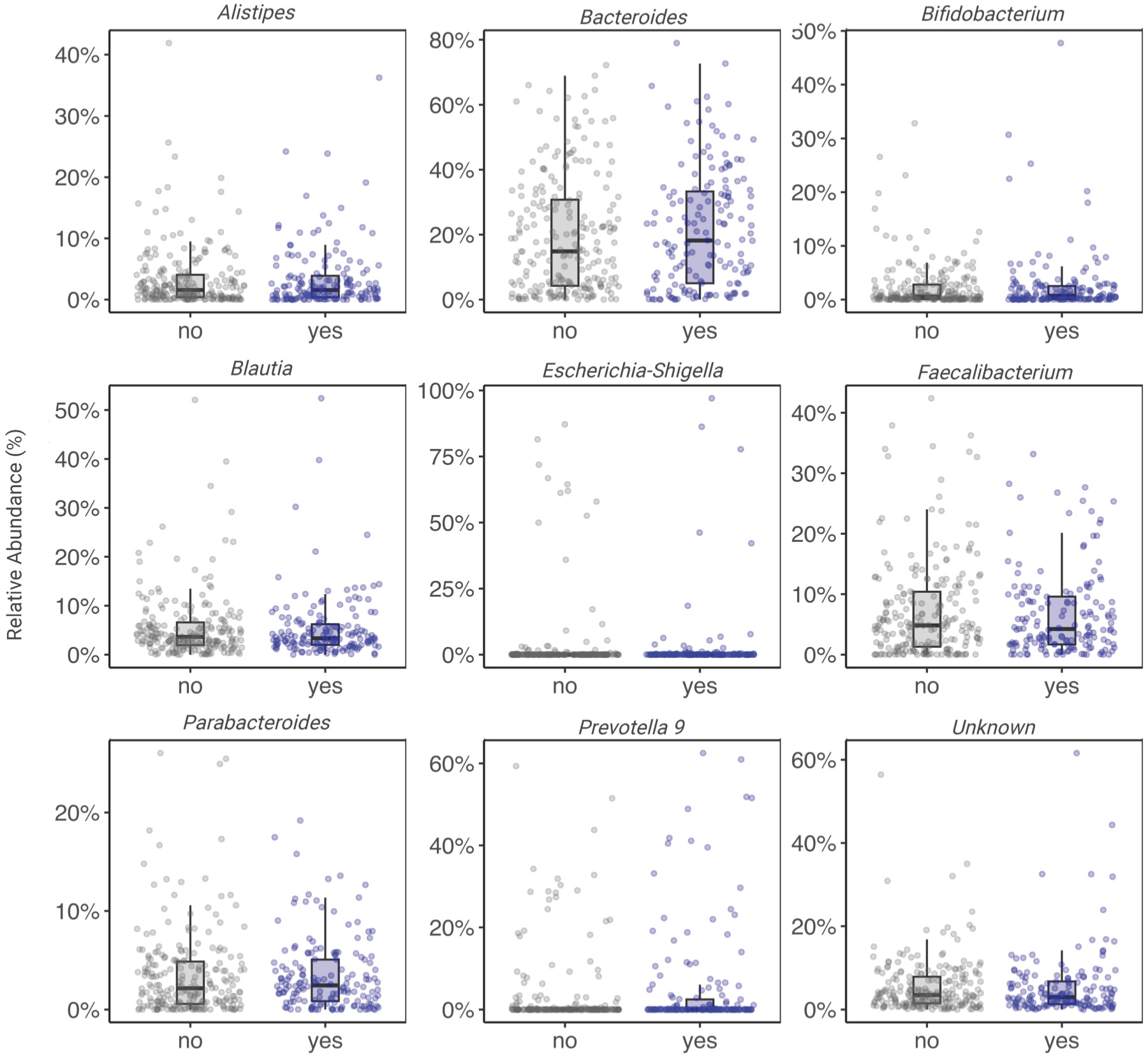
Most abundant genera for Veterans in the United States-Veteran Microbiome Project by lifetime history of homelessness.

Microbial community structures from weighted UniFrac were significantly different between Veterans with and without a lifetime history of homelessness after adjustment for age, gender and education level (**Fig 2A**, PERMANOVA, *p* = 0.047) and approached statistical significance for Unweighted UniFrac (**Fig 2B**, PERMANOVA, *p* = 0.086). No interactions between lifetime history of homelessness and covariates were significant. The communities were not significantly different in Bray-Curtis analysis. Significantly differentially abundant taxa of Veterans with a lifetime history of homelessness, relative to Veterans without a lifetime history of homelessness, after adjustment for age, gender, and education level included *Akkermansia* (*p* = 0.014), *Acidaminococcus* (*p* = 0.023), *Barnesiella* (*p* = 0.038), and *Ruminococcaceae* UCG-003 (*p* = 0.041) (**Table S3**). The significance was not retained after adjustment for multiple tests. Moreover, *Akkermansia* was observed at a lower relative abundance in Veterans that reported a lifetime history of homelessness (1.07 ± 3.85) compared to those that had no homeless experiences (2.02 ± 5.36). In contrast, *Prevotella 9* was observed at lower relative abundance for Veterans with no history of homelessness (3.36 ± 9.46) compared to Veterans with a lifetime history of homelessness (5.40 ± 12.8).

**Fig 2.**
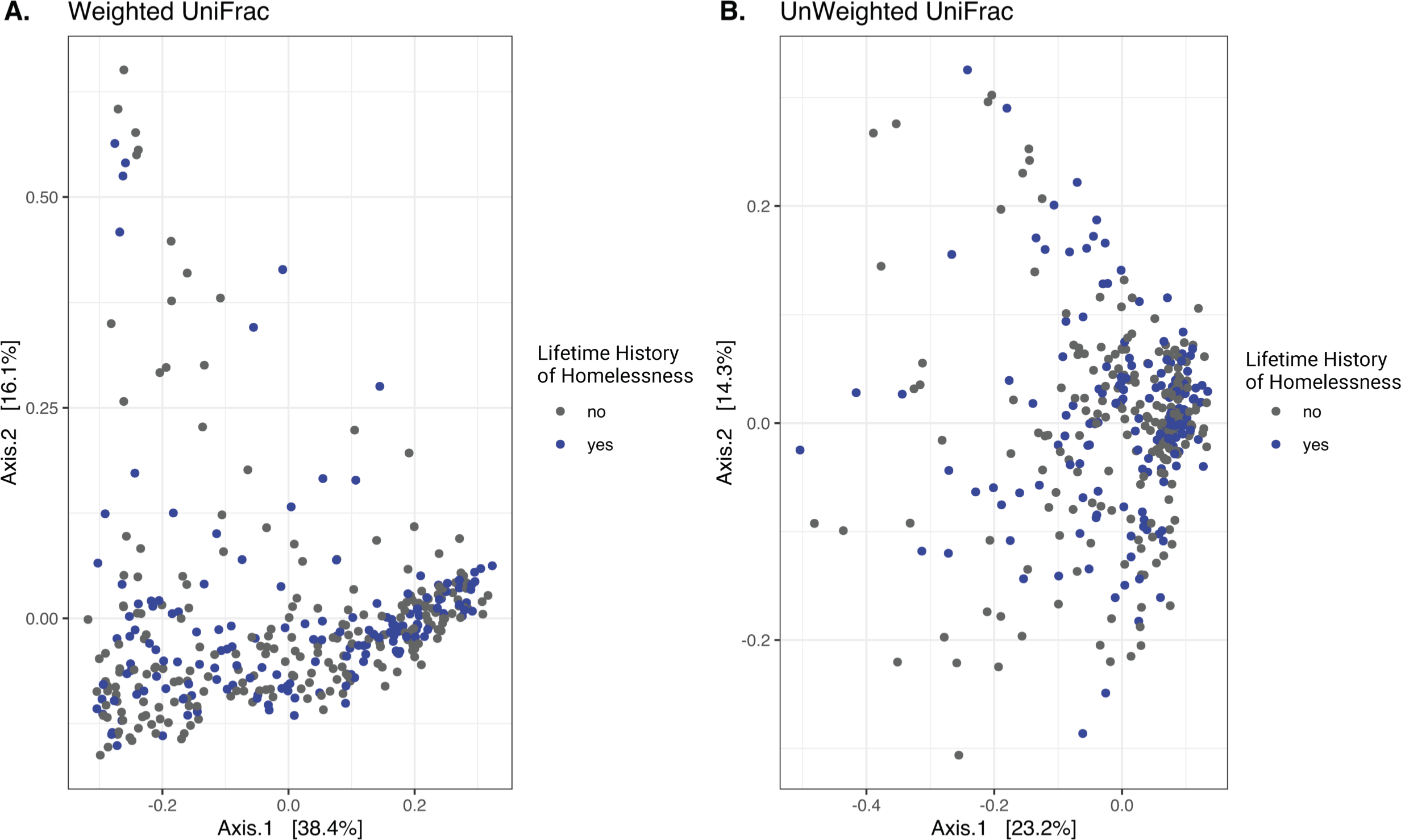
Principal Component Analysis Results of Veterans with and without a lifetime history of homelessness for **(A)** Weighted UniFrac and **(B)** Unweighted UniFrac.

For Veterans with a lifetime history of homelessness, the most highly associated external factors from Weighted UniFrac were education level, and SF-36 MCS scores (**Fig 3A, Table S4**). In Veterans with no history of homelessness, the only significant external factor for Weighted UniFrac was gender. Military sexual harassment (MST 1) and assault (MST 2) had greater effect sizes for Veterans that never experienced homelessness compared to individuals with a lifetime history of homelessness. While military sexual harassment and assault were more likely to be reported by females compared to males (MST 1 and MST 2, χ^2^ *p* < 0.0001), when isolated neither male nor female Weighted UniFrac was significantly different by either category of military sexual trauma (i.e., harassment or assault).

**Fig 3.**
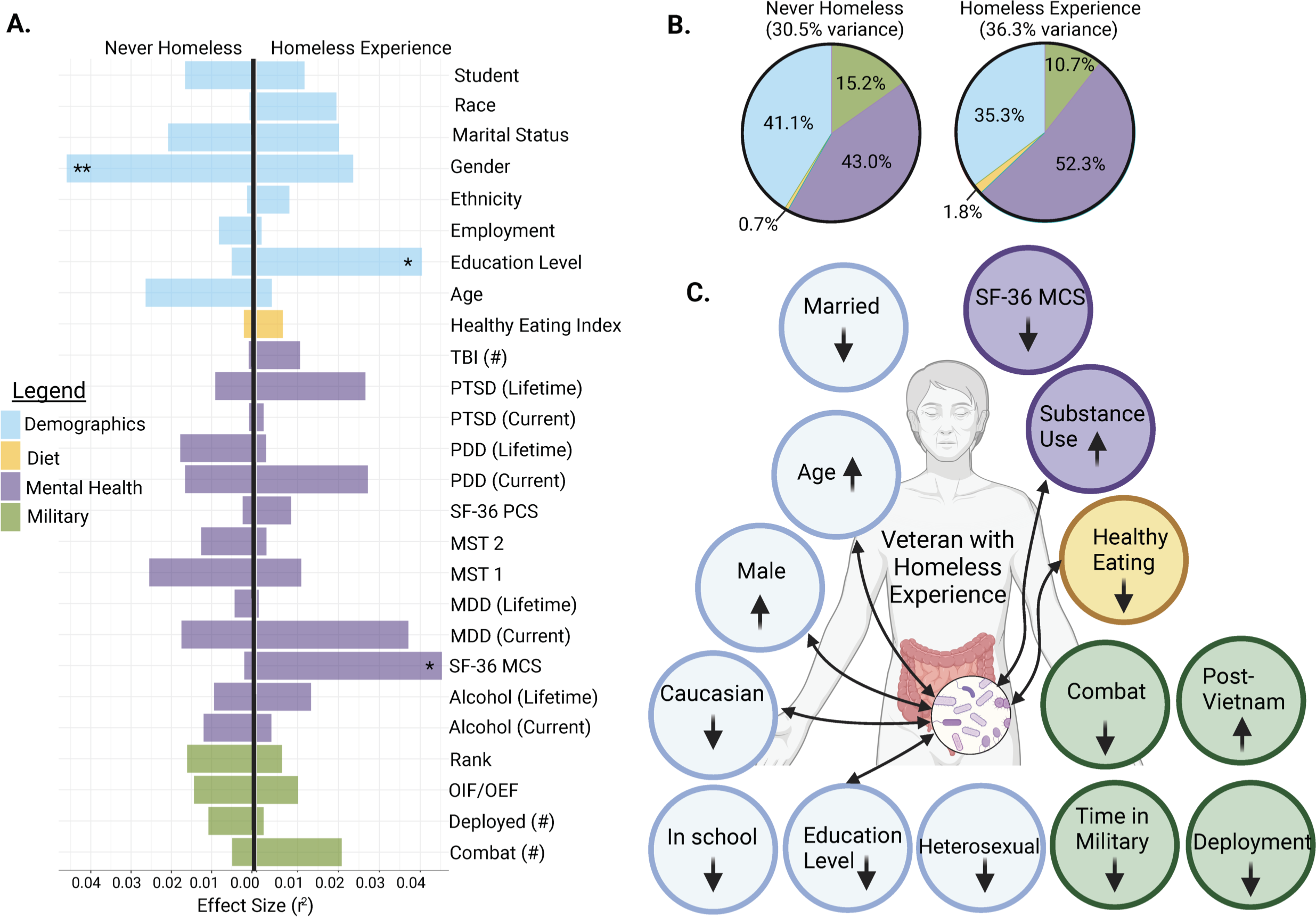
Weighted UniFrac Distance Metric between Veterans with a lifetime history of homelessness and Veterans with no history of homelessness for **(A)** metadata categories effect sizes that were most influential to CCA through PERMANOVA analysis (* *p* < 0.05, ** *p* < 0.01), (B) variance explained for the main groupings in part A, **(C)** statistically different external factors (arrows within circles indicate direction of difference and arrows between circles show influences of factors on microbiome from past studies). Overall colors: blue for demographics, yellow for diet, purple for mental health measures, and green for military-related categories.

Overall, the twenty-six metadata categories tested accounted for 30.5% of the variance explained in the never homeless group and 36.5% in those that had experienced homelessness (**Fig 3B**). Having a history of homelessness increased the effect size for microbiota community influences in the broad categories of mental health measures (52.3% vs. 43.0%) and diet (1.8% vs. 0.7%) and decreased the effect size in demographics (35.3% vs. 41.1%) and military-related factors (10.7% vs. 15.2%). Significant demographic differences were observed for the categories of gender (lifetime history of homelessness, 90.9% male vs. 75.9% female, χ^2^ *p* = 0.026), age (lifetime history of homelessness mean 51.9 years old vs. 45.9, *t*-test *p* < 0.001), and education level at bachelor’s degree or higher (lifetime history of homelessness 23.0% vs. 47.0%, *t*-test *p* < 0.001) (**Table S1 &** **Fig 3A**).

### Fecal Microbiome - Homelessness: Episodes/Current

To further delineate the impact of homelessness, Veterans were grouped into categories based on number of reported experiences of homelessness: no lifetime history (*n* = 228); one experience (*n* = 65); two to three experiences (*n* = 58); and, four or more experiences (*n* = 37). The delineation did not result in significant differences for alpha diversity, beta diversity, or the five most abundant taxa (**Figures S2 & S3**). Differential abundant taxa prior to adjustment included *Akkermansia, Bifidobacterium*, *Corynebacterium 1*, *Collinsella*, *Mogibacterium*, and *Slackia* (**Table S5**). No taxa were significantly different after correction for multiple testing.

In addition, 31 of our 388 participants (8.0%) in the present study reported being currently homeless. The two groups of currently homeless and not currently homeless had similar alpha diversity (e.g., Shannon Diversity Index, Pielou’s Evenness, and Observed ASVs) and overall microbial communities (Weighted UniFrac, Unweighted UniFrac, Bray-Curtis) (**Fig S4 & S5**). There were no statistical differences between the two groups at the genus level (**Table S6).** CCA for Weighted UniFrac revealed the most highly associated metadata to current homelessness based on effect size (*r^2^*) was lifetime alcohol use disorder, education level, current major depressive disorder (MDD), and SF-36 MCS scores (**Table S7**). Differences in the cohort data between currently homeless and no reported history of homelessness were statistically significant for lifetime alcohol use disorder (χ^2^*p* < 0.0001), education level (Fisher’s exact test *p* = 0.0005), and Healthy Eating Index score (Independent samples *t*-test, *p* = 0.008).

## Discussion

This is the first study aimed at comparing the gut microbiome of those with a lifetime history of homelessness or current homelessness to those without such a history. We also explored how dietary quality and military-related factors (e.g., combat history) may have contributed to differences in outcomes (e.g., demographic, microbiome) among members of this cohort. Our findings highlight that diet- and microbiome-related differences between cohorts might be actually rooted in psychosocial factors, with origins that mainly pre-dated military service (e.g., social determinants of health; see **Fig 2C**).

In terms of diet, Veterans on average scored higher than the national average on the HEI, indicating higher adherence to dietary guidelines. This was expected based on earlier findings from this team (51). However, Veterans with and without a history of homelessness significantly differed on HEI total scores and several sub-scores. While scores were similar on dietary habits pertaining to pro-commensal fiber intake (e.g., Total Vegetables, Beans & Greens), those with no reported history of homelessness had higher-quality diets than those who reported a lifetime history of homelessness, particularly in regard to intake of Fruit, Plant proteins, and Whole grains. It should be noted that participants with a lifetime or current history of homelessness on average consume *fewer* Added Sugars compared to those reporting no history of homelessness, which is a habit associated with better dietary quality. Given the larger context of dietary patterns, however, the evidence suggests current and/or historical experiences of homelessness are associated with a poorer overall diet.

This is consistent with previous literature regarding dietary quality among people who are unhoused/have unstable housing, which suggests that poorer access to food, inadequate food storage resources, and a lack of cooking amenities can all contribute to poorer dietary quality (52). Moreover, substance and alcohol use disorders, both of which were significantly more prevalent among Veterans who reported a lifetime or current history of homelessness, have been found to be independently associated with poorer quality diets (53), potentially further explaining our findings.

In comparing gut microbial communities between those with a lifetime history of homelessness versus those without this history, differential abundance analysis revealed those without a homeless experience had higher abundance of *Akkermansia* and lower *Prevotella 9.* Interestingly, Graner and colleagues found that when compared to those without alcohol-related liver disease (ALD) those with ALD exhibited decreased relative abundance of *Akkermansia muciniphila*, an intestinal commensal that promotes barrier function in part by enhancing mucus production (54). Recent interventions have used probiotic supplementation of *Akkermansia* for several health outcomes (55, 56), and it might have potential for those with a history of homelessness; however further research is required. While the observed *Akkermansia* trend was as expected, finding lower *Prevotella 9* in Veterans with no history of homelessness was unexpected. This genus has been reportedly associated with high fiber, low-fat diets (57, 58) with beneficial effect to improve glucose metabolism (59). While lower, *Prevotella 9* was not significantly different between the two groups (ancom-bc2). The observed trend would be interesting to explore with additional sequencing resolution (e.g., whole genome sequencing) or evaluation of short-chain fatty acid concentrations.

On the whole, influences on the gut microbiota are multi-factorial, and Veterans included in this study with a history of homelessness had histories of complex mental and physical health conditions. Generally, we observed that the gut microbiome of Veterans with a lifetime history of homelessness was driven more by their mental health conditions relative to Veterans that had never been homeless. For example, significantly lower perceived mental health functioning (SF-36 MCS) was noted among those with a history of homelessness when compared to those without such a history, and lower scores were also found to be associated with differences in gut microbial composition for those with a homeless experience. Interestingly, history of current and/or lifetime alcohol use disorder was significantly different between the two groups, but not identified as a driver of gut microbiota composition. This was a surprising discovery considering the known influences of alcohol on the microbiota (60, 61).

Diet and homelessness appear to both be contributing factors that alter the gut microbiome, yet it remains unclear if either element is directly causative to gut microbiome community composition. Indeed, many of the significant factors between Veterans with a lifetime history of homelessness and those who have not experienced homelessness (**Fig 2C**) have been shown to influence the gut microbiota among cohorts without a history of military service. For example, the impact of aging on altering the gut microbiome is well documented (62–64), and parallel research in advanced aging has shown gut microbiome modulating effects such as reduction in intestinal barrier function and increased systemic inflammation (65). Other external factors that were significantly different for Veterans that had experienced homelessness included increased prevalence of alcohol use disorder and substance use disorder, both factors that also have been associated with microbiome dysbiosis (61, 66). Similar to aging, alcohol use disorder and substance use disorder reduce intestinal barrier function and increase inflammation (67). Taken together, the differences in Veterans that have experienced homelessness could be similarly influencing the gut microbiota, and therefore could alter the microbiota in a non-linear manner.

Among this cohort of Veterans, initial efforts were aimed at identifying potential associations between military-related factors, homelessness, and the gut microbiome. Our results suggest that differences may have been more related to psychosocial stressors that pre-dated military service. Notably, among those with a lifetime history of homelessness the education levels were lower. Access to education, is one of many, often co-occurring social determinants of health known to impact health disparities (68). Fielding and colleagues suggested that health, which is not simply the absence of disease, should be conceptualized within broader contexts (e.g., cultural factors, social and community networks), as well as over the course of a lifetime (69). Further support for this life course perspective included findings suggesting that the number of episodes of homelessness did not appear to impact microbiome findings. In addition, there were microbial community differences for those that were noted as currently homeless versus those with a lifetime history of homelessness. Interestingly, Veterans that had experienced homelessness had less time in the military, along with fewer deployments and combat experiences, compared to Veterans that had never been homeless. Possibly this is due to a previously described “healthy deployer effect” (70, 71), a theory that hypothesizes deployers tend to be healthier than non-deployers.

This study had several limitations. Some of the sub-cohort sample sizes were small (e.g., currently homeless Veterans, women Veterans); thereby contributing to a lack of clarity to the etiology of some of the findings (e.g., history of sexual trauma). Larger longitudinal efforts are indicated in which women Veterans with and without a history of homelessness are included.

The initial study was not designed to evaluate homelessness and therefore did not include a standardized definition of homelessness, such as the standard VA screening items for housing instability and homelessness. The study also did not evaluate the time since homelessness (the point in time at which a given Veteran could officially be classified as housed), or duration of individual experiences of homelessness. A better understanding of those factors might enable a more accurate understanding regarding associations between homelessness and fecal microbiome composition, as well as lead to treatment options for conditions frequently observed among those who are homeless. In contrast, a strength of this paper is that it presents the first study of fecal microbiome diversity and community composition among those who are homeless and outlines relationships between the microbiome, demographics dietary data, and mental health conditions among this at-risk cohort.

## Conclusions

History of homelessness was associated with composition of the gut microbiome, although these associations may be due to other factors that are also associated with homelessness. Diet and gut microbiome differences noted among Veterans with a lifetime history of homelessness may be more related to broader psychosocial factors that pre-dated individuals’ history of military service. Increased efforts are required to longitudinally evaluate the complex relationships between homelessness, the gut microbiome, and mental and physical health outcomes.

## Acknowledgements

This project was in part supported by the Department of Veterans Affairs (VA) Rocky Mountain Mental Illness Research Education and Clinical Center (MIRECC) for Suicide Prevention. The views, opinions, and/or findings contained in this article are those of the author(s) and should not be construed as an official Department of Defense or VA position, policy, or decision unless so designated by other documentation. All individuals equally contributed to this manuscript.

## Data Availability

Demultiplexed single-end sequences were deposited in the NCBI Sequence Read Archive (BioProject accession ID: PRJNA1010779).

